# Efficient calculation of fluid transport in porous media with moving boundaries

**DOI:** 10.64898/2025.12.15.694505

**Authors:** Huabing Li, Wenjie Lu, Badr Kaoui, James W. Baish, Lance L. Munn

## Abstract

A novel hybrid model combining the lattice Boltzmann (LB) and finite difference (FD) methods is proposed to simulate transport in through a junction of actively contracting lymphatic vessels, while also handling flow of interstitial liquid in the surrounding porous tissue. Details of the dynamically flexing walls and valves in the lymphatic vessel and its near vicinity are modeled using a high-resolution LB method, whereas overall efficiency was significantly improved by using low-resolution FD in the larger tissue domain distant from the vessel. Pressure and velocity conditions at the interface between subdomains of the two numerical methods are matched by imposing a partial bounce-back ratio in LB corresponding to the permeability coefficient *κ* in Darcy’s law for flow through porous media. Parameters governing the match between the algorithms at their interface can be estimated from the Kozeny-Carman relationship for porous media and further refined with a simpler, parallel flow geometry that also serves to validate the method. Test calculations show that the hybrid method is roughly four times faster than the LB method and permits computation over significantly larger domains. This method should be applicable to a large range of problems involving fluid flow in porous media with embedded conduits that have non-stationary boundaries.

**Author summary:** It is generally acknowledged that the finite difference method (FDM) is faster and requires less memory than the lattice Boltzmann method (LBM) for comparable domain sizes. However, LBM performs better for simulating fluid flow near complex, deformable, or moving boundaries. For this reason, it can be beneficial to create hybrid models that combine FDM and LBM. In this work, we use such a hybrid model to simulate a contracting lymphatic bifurcation in fluid. Our goal is to demonstrate the model’s robustness and high efficiency.

## Introduction

The LBM is a numerical method suitable for computing flow in complex geometries. For example, flows with moving boundaries [1], including through contracting channels with elastic valves such as those in lymph vessels, have been readily solved [2, 3].

However, the memory and computational costs of LBM become prohibitive when only a few complicated features are embedded in a much larger domain governed by simple flow through a porous continuum. Instead, the FDM is better suited to the task of computing flow in a porous domain governed by Darcy’s law. The computational and memory costs of the LBM are ∼ 4 times more than for FDM algorithms for the same domain size calculated for 200000 steps [4, 5]. Compared with mesh refinement with LBM, FDM is more efficient for simple flow through the porous continuum.

In the case of fluid drainage in biological tissues, extravascular tissues constitute a porous medium, but embedded lymphatic vessels undergo cyclic contractions, making the fluid dynamics more complex. In addition to fluid flow in tissues, there are other physical and biological systems that include fluid flow through porous media and moving boundaries. For example, in cases where resources percolate through geological substrates and other industrial applications, such as tissue scaffolds used in bioengineering, modern emerging battery technologies (for example, flexible smart-phones), nutrient delivery from blood vessels, flow in irrigation systems, and oil recovery operations. Modeling large domains is a challenge for the LBM because of computational costs. On the other hand, FDM methods can be applied to large domains, but are not easily adapted to moving boundaries. Thus, a hybrid approach may be more efficient for many applications.

## Materials and methods

### Formulation in Respective Domains

Darcy’s law is used to model flow in the porous region outside the vessel wall, which includes all of the FDM domain and some of the LBM domain (Fig. 4) [6]:

**Fig 1.**
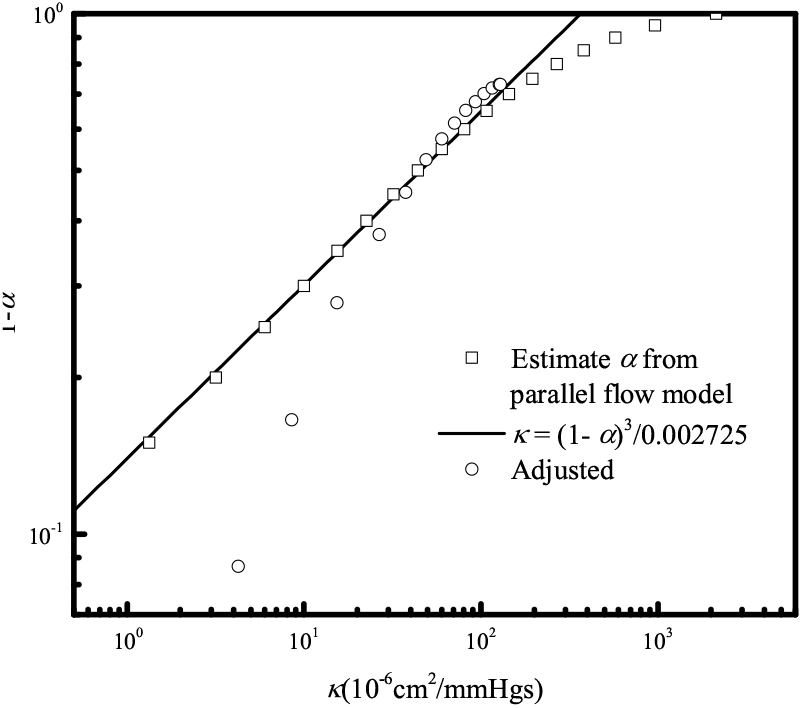
Mapping of the bounce-back ratio *α* to the permeability *κ* of porous media. Data presented with circles ∘ are adjusted from the magnitude of the fluid velocity map.

**Fig 2.**
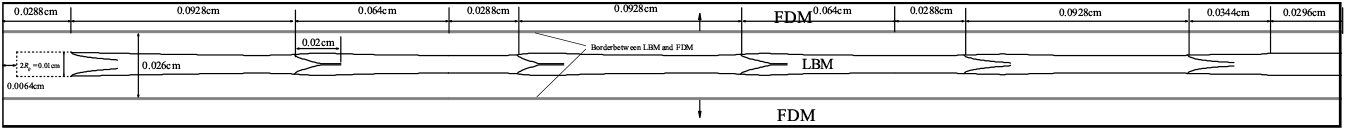
Schematic diagram of the hybrid model. Moving the borders upward and downward at same time can change the areas of FDM and LBM, but the total area is not changed.

**Fig 3.**
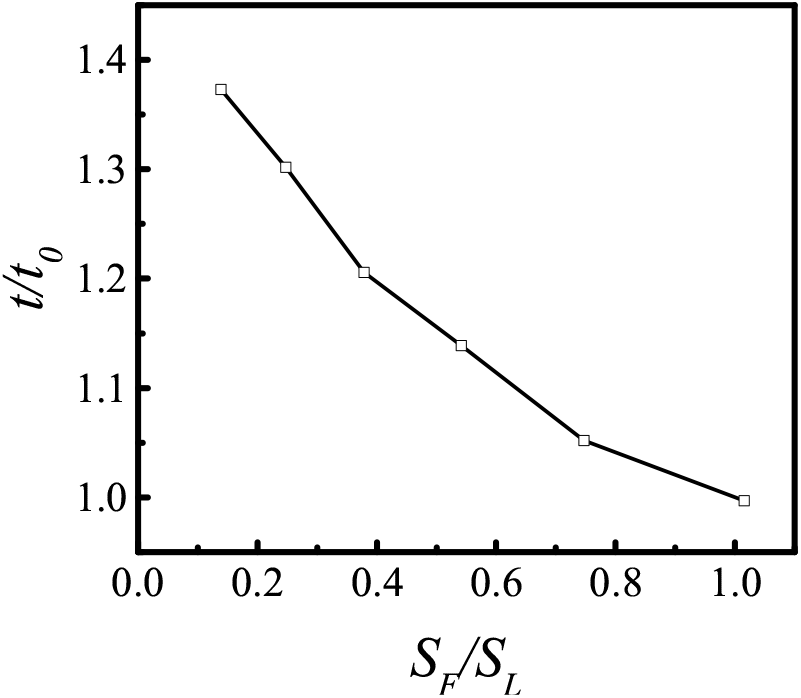
Dimensionless calculation time for a single vessel with three lymphangions (six valves) contracting to drive flow. The central area, *S*_*L*_, is calculated using LBM. The upper and lower subdomains are calculated using FDM. The total area is *S*_*F*_ . *t*_0_ is the calculation time when *S*_*F*_ */S*_*L*_ = 1.

**Fig 4.**
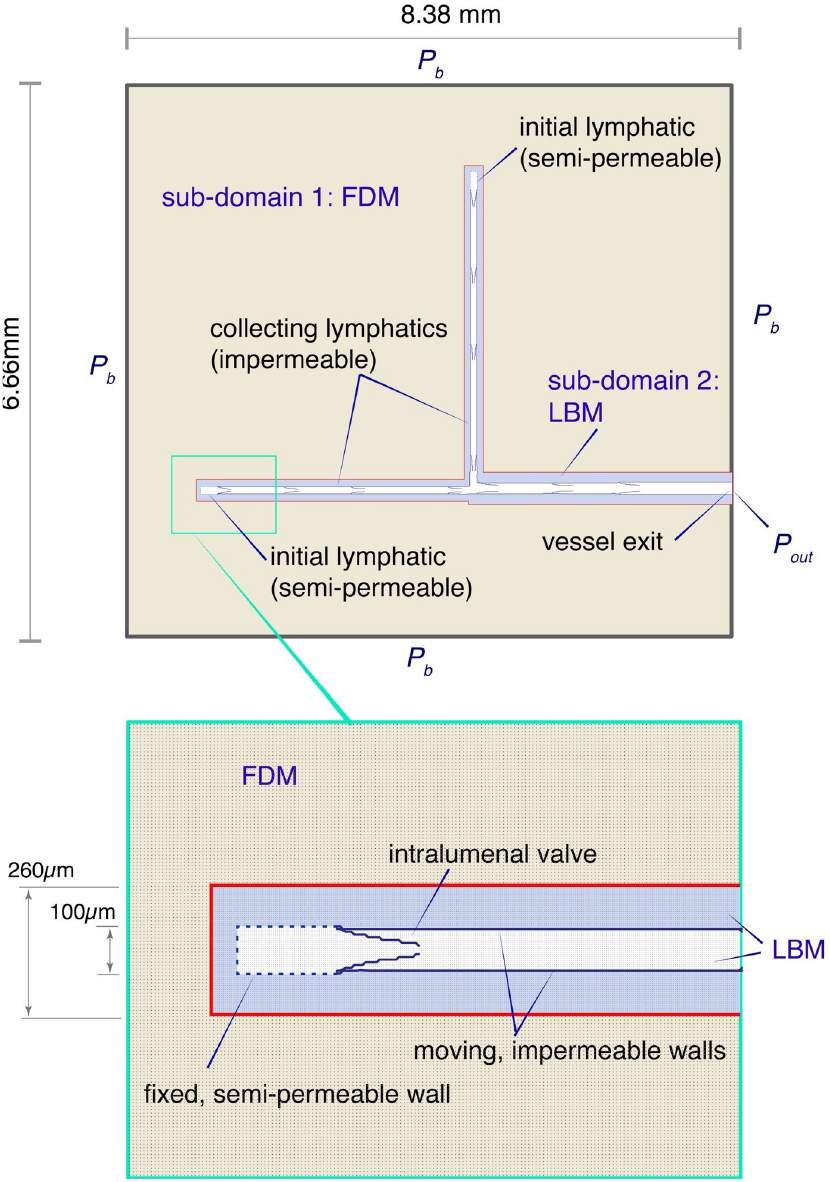
Schematic diagram of computational domains. The outer sub-domain (tan) is handled by FDM, while the blue sub-domain is handled by LBM. The red line is the numerical interface. The pressures at the outer domain boundary (*P*_*b*_) and the vessel outlet (*P*_*out*_) are specified.

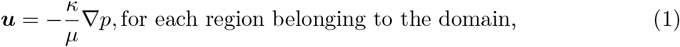

where ***u*** is the fluid velocity, *µ* is the dynamic viscosity of the fluid, *p* is the pressure, *κ* is the tissue solvent permeability. The pressure distribution can be evaluated from the continuity equation:

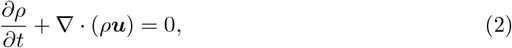

where *ρ* is the density of the fluid. The regions of the LBM domain that lie outside the vessel boundary contain porous media and are also solved using Darcy’s law. As an alternative to solving directly for the pressure, as is typically done with the FDM, we anticipate the need to link the FDM result to the LBM method, where the density and pressure are related via the state equation [7]

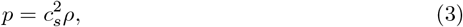

where *c*_*s*_ is the speed of sound. In the FDM domain, the central difference of Eq. (2) is

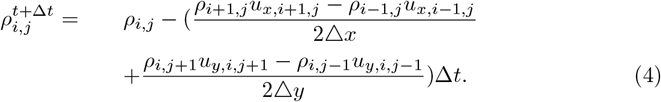

where Δ*x* = Δ*y*, and Δ*t* are the grid spacing on the lattice and time step, respectively. Velocity components are then recovered from the difference form of Eq. (1) where we have incorporated Eq. (3)

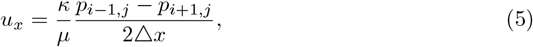

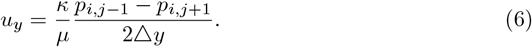

We update the pressures at each time step using Eqs. (3) and (4) and the fluid velocities using Eqs. (5) and (6).

Computations within the LBM domain proceed differently. LBM recovers the solutions of the full Navier-Stokes equations (in the limit of small Mach number and Knudsen number) [7], which is required inside and near the moving walls of the vessel but is unnecessarily complex in most porous domains. The BGK LBM assumes a weakly compressible fluid [7].

The LBM has proven effective in simulating fluid-structure interactions. This can be achieved either by coupling it with the immersed boundary method [8] or by using it independently by implementing special bounce-back boundary conditions on the surface of a structure [1–3]. The latter approach requires careful computation of momentum exchange at the structure surface, which may feature complex and curved geometries.

Additionally, this approach can ensure the impermeability of the fluid at the fluid-structure interface. This approach can be extended to semi-permeable boundaries by imposing a partial bounce-back condition, where some fraction of the fluid is allowed to pass through the wall, while the rest is reflected [9]. This approach can be used to model fluid leakage across the boundary [10]]. By imposing partial bounce-back to each point in the domain, we can also simulate flow through porous media [2]. This approach is used to model the porous medium within the LBM sub-domain located outside the vessel (Fig. 4, blue region). In this region, a portion of the streamed distribution function is reflected backward at every point in the sub-domain, following Eq. (7), which is a modified form of the well-established bounce-back boundary condition. Increasing this reflected fraction effectively reduces the physical permeability in accordance with Darcy’s equation.

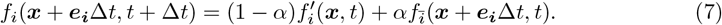

and,

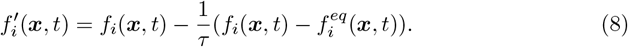

where *f*_*i*_ is the distribution function. 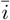 indicates the reverse direction of *i*. ***e***_***i***_ is the discrete velocity along the *i* direction, and *α* is the bounce-back ratio. 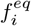 is the equilibrium distribution function:

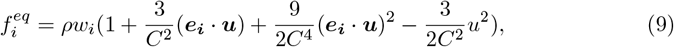

here, *C* = Δ*x*/Δ*t* is the lattice’s migration speed.

*w*_0_ = 4/9, *w*_1,2,3,4_ = 1/9, *w*_5,6,7,8_ = 1/36 for the D2Q9 lattice used in the present study.

### Coordinating calculations between the LBM and FDM domains

A challenge with the domain in Fig. 4 is to transfer information between the LBM and FDM sub-domains at each time step. To do this, we also use the equilibrium distribution function to adjust the boundary distribution function on the LBM side of the interface between the two methods, but *ρ* and ***u*** are transferred from the FDM domain before the streaming step of LBM. Before we update the FDM domain using Eqs, (4) -(6), we transfer the new velocity of the boundary of the LBM domain, but the pressure on the FDM side on the boundary must be extrapolated from the pressure of the previous step of the FDM domain.

On the LB side of the bridging interface, the distribution functions required for the LB streaming step are initialized as equilibrium distribution functions, with the density and velocity computed using the FDM. The velocity obtained from LB is then applied on the FD side of the interface in Eq. (4). However, the pressure is not taken from LB; instead, it is extrapolated from the pressures computed by FD at the nodes closest to the interface.

For the outer boundary condition of the entire domain, we use constant pressure *p*_*b*_. Velocities at the boundary are extrapolated from the FDM domain. A pressure boundary condition Pout [11] is applied at the outlet of the vessel where flow exits at the right boundary of the LBM domain (Fig. 4). For computational efficiency and to decrease memory requirements, we need to consider the location of the bridging interface to minimize the number of computational nodes within the LB subdomain. The primary criterion in this study is to ensure that sufficient lateral space is provided for the vessel wall to move freely, without impinging on the bridging interface.

### Calibration of the LBM bounce-back parameter

We first specify the permeability of the FDM sub-domain according to known tissue porosities [12]. For the small region of porous medium near the vessel and inside the LBM subdomain, Darcy’s law is approximated with LBM by applying a partial bounce-back condition within the tissue. To estimate the LBM bounce-back ratio needed to reproduce the permeability in the adjacent tissue (calculated with FDM), we constructed a simplified rectangular domain (see supplementary information Fig. S1). The same pressure gradient was applied across the domain for calculating flow using LBM or FDM. We then iteratively adjusted the bounce-back ratio *α* in LBM to match the flow calculated by FDM. Increasing *α* in the porous LBM domain reduces the effective value of *κ*, much like reducing the porosity *ε* of an actual medium reduces *κ* (Fig. 1). For example, the Kozeny-Carman relationship [13] for a packed bed of particles predicts *κ ∼ ε*^3^/(1 *− ε*)^2^, where if we replace *ε* with 1 *− α* closely matches our observation that *κ ∼* (1 *− α*)^3^*/α*^2^ for the low values of *κ* of most practical interest. For example, the maximum permeability in the mouse tail skin is about 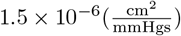 [14], corresponding to an approximate *α* = 0.9663. In practice, because the LBM permeability is very sensitive to *α*, fine adjustment may be necessary for specific applications to optimally smooth the velocity across the LBM/FDM boundary (see supplementary information, Fig. S2).

### Calibration of the calculation efficiency of the hybrid model

To compare the calculation efficiencies of the LBM and FDM approaches, we consider only the left branch of Fig. 4, but extend the length so that it consists of three lymphangions, defined by six valves (Fig. 2). Flow in and near the central vessel is calculated by LBM, while flow in the porous media at top and bottom is calculated by FDM. The lattice size is the same for the two regions. We then expand the size of the central subdomain by moving borders upward and downward at same time; in this way, the total area was not changed, only the relative domain areas calculated by LBM vs. FDM. The results show that enlarging the area calculated by FDM reduces calculation time (Fig. 3), consistent with our preconception that FDM is more efficient than LBM. But the hybrid model wants to synchronize the two methods, thus the calculation speed of FDM will be lower than unmixed FDM. Furthermore, the relationship in Fig. 3 is nonlinear because FDM requires less memory, and data transfer from one segment to another by MPI takes less time.

## Results

Here, we demonstrate the implementation of a hybrid FDM - LBM approach to model fluid flow in a porous biological tissue that contains contracting lymphatic vessels. This approach retains the flexibility of LBM in a small sub-domain in and near contracting vessels while using the simpler and faster FDM in the surrounding porous tissue.

Fig. 1 illustrates the entire computational domain, which is divided into two subdomains. Specification of the boundary between the two sub-domains is somewhat arbitrary, but can affect overall computational efficiency. The LBM domain should be large enough to encompass sufficient nodes around the moving boundaries so that the complex fluid dynamics in these regions are captured. On the other hand, minimizing the size of this subdomain increases efficiency. Within each sub-domain, the physical quantities are computed independently using either FD or LB. The physical quantities are then exchanged between the sub-domains across the bridging interface, represented by the red border in Fig. 1.

In the porous region outside the vessels, both methods solve Darcy’s law. The numerical challenge for this hybrid LBM - FDM approach is to match pressure and velocity conditions at the boundary between their respective sub-domains within the porous region.

### Demonstration of the method: Lymph transport from tissue by pumping lymphatic vessels

To demonstrate the method, we simulate fluid drainage from tissue, driven by an imposed boundary pressure and active contractions of the lymphatic vessels, which have intralumenal valves (Fig. 4). Fluid enters at the boundary, convects through the porous media of the FDM domain, crosses the subdomain boundary into the porous media of the LBM domain, and then enter into the two lymphatic vessels at the initial lymphatic segments at top and bottom left of the domain. The collecting vessel walls periodically contract due to forces imposed by vascular muscle cells, as described previously [15].

The contractions produce flow through the vessels, but also lower the pressure outside the vessel wall, which propagates through the domain (see Fig. 5 (A-C)). Because of the careful matching of subdomain permeabilities by calibrating *α*, smooth continuity is achieved at the boundary between the FDM and LBM subdomains.

**Fig 5.**
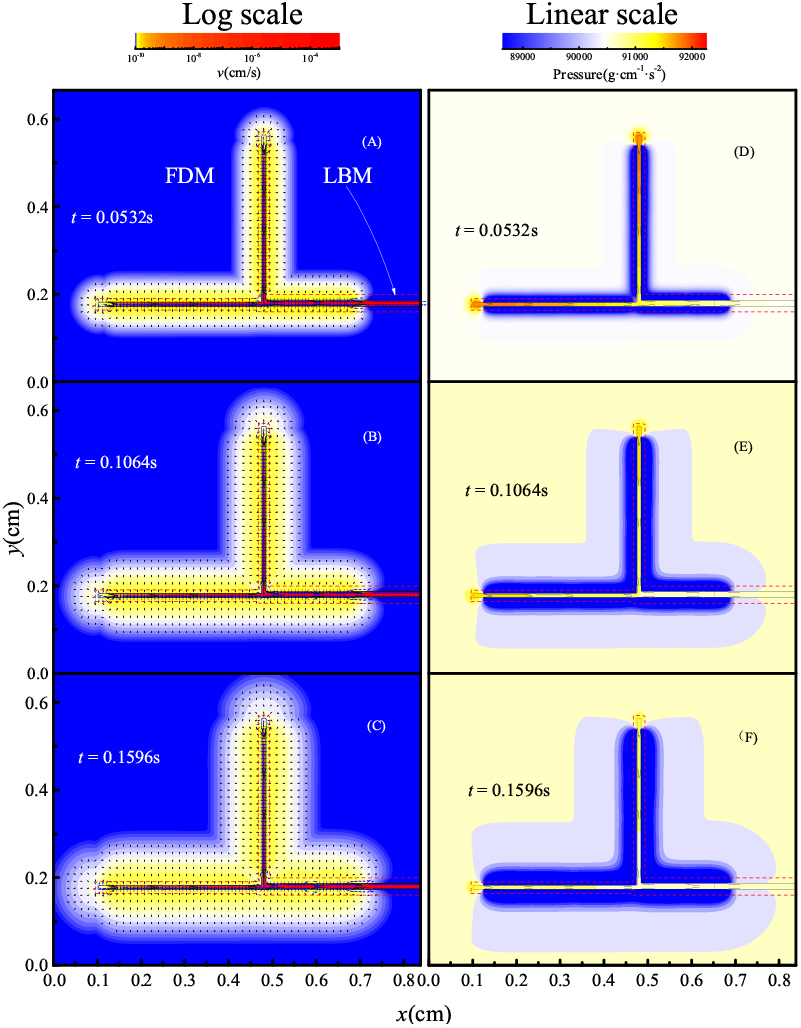
Velocity field and pressure dis tFir ib.0ution in the tissue. In panels A - C, the amplitude of velocity is logarithmic; the corresponding pressures are shown in panels D - F. *α* is modified by iteration.

## Discussion

Our numerical hybrid method is well-suited for predicting fluid flow in porous media with complex networks of flexible ducts. At this stage of development, we have not accounted for changes in porosity due to boundary motion and tissue deformation, which may have a significant impact [16]. However, these effects could, in principle, be easily incorporated into our hybrid method.

## Conclusion

We have presented a hybrid, multi-scale approach to match hydrodynamic quantities computed in different regions by partial bounce back of LBM and Darcy’s law with FDM. The method leverages the computational efficiency of FDM while providing the ability to include complex, moving boundaries using the LB method. The potential impact of using such a hybrid approach is underscored in our comparison of computational efficiencies for the two methods (Supplementary Information). Using a simple rectangular domain and equivalent pressure gradients, the FDM is ∼ 4 times faster than LBM (see Fig. S1 in supplementary information calculated by LBM and FDM respectively). But in the hybrid model, the calculation speed depends on the slowest method, because the data wants to be synopsized on each step. The separate FDM and LBM domains are matched by adjusting the LBM bounce-back parameter. Initially estimated using a simple geometry with no moving boundaries, fine adjustment can be made in the actual domain to minimize discontinuities.

## Acknowledgments

This work was supported by NIH R01 CA284603 and U01CA261842 (LLM) and ANR (ANR-20-CE45-0008-01) (BK).

## Notes

### Competing Interest Statement

The authors have declared no competing interest.

## References

1. Aidun CK, Clausen JR. Lattice-Boltzmann method for complex flows. Annual review of fluid mechanics. 2010;42(1):439–72.

2. Li H, Zhang J, Padera TP, Baish JW, Munn LL. Fluid dynamics and leukocyte transit in the lymphatic system. PNAS nexus. 2024;3(6):pgae195.

3. Li H, Wei H, Padera TP, Baish JW, Munn LL. Computational simulations of the effects of gravity on lymphatic transport. PNAS nexus. 2022;1(5):pgac237.

4. Yoshino M, Matsuda Y, Shao C. Comparison of accuracy and efficiency between the lattice Boltzmann method and the finite difference method in viscous/thermal fluid flows. International Journal of Computational Fluid Dynamics. 2004;18(4):333–45.

5. Wichmann KR, Kronbichler M, Löhner R, Wall WA. A runtime based comparison of highly tuned lattice Boltzmann and finite difference solvers. The International Journal of High Performance Computing Applications. 2021;35(4):370–90.

6. Hubbert MK. Darcy’s law and the field equations of the flow of underground fluids. Transactions of the AIME. 1956;207(01):222–39.

7. Sukop M, DT Thorne J. Lattice Boltzmann Modeling Lattice Boltzmann Modeling. Springer; 2006.

8. Bielinski C, Xia L, Helbecque G, Kaoui B. Mass transfer from a sheared spherical rigid capsule. Physics of Fluids. 2022;34(3).

9. Succi S. The Lattice Boltzmann Equation for Fluid Dynamics and Beyond. Oxford University Press; 2001.

10. Zhao Yl, Wang Zm. Multi-scale analysis on coal permeability using the lattice Boltzmann method. Journal of Petroleum Science and Engineering. 2019;174:1269–78.

11. Zou Q, He X. On pressure and velocity boundary conditions for the lattice Boltzmann BGK model. Physics of fluids. 1997;9(6):1591–8.

12. Swartz MA, Fleury ME. Interstitial flow and its effects in soft tissues. The Annual Review of Biomedical Engineering. 2007;9(1):229–56.

13. Childs EC, Collis-George N. The permeability of porous materials. Proceedings of the royal society of London series a mathematical and physical sciences. 1950;201(1066):392–405.

14. Swartz MA, Kaipainen A, Netti PA, Brekken C, Boucher Y, Grodzinsky AJ, et al. Mechanics of interstitial-lymphatic fluid transport: theoretical foundation and experimental validation. Journal of biomechanics. 1999;32(12):1297–307.

15. Kunert C, Baish JW, Liao S, Padera TP, Munn LL. Mechanobiological oscillators control lymph flow. Proceedings of the National Academy of Sciences. 2015;112(35):10938–43.

16. Lai W, Mow VC, Roth V. Effects of nonlinear strain-dependent permeability and rate of compression on the stress behavior of articular cartilage; 1981.

